# Retention time and fragmentation predictors increase confidence in variant peptide identification

**DOI:** 10.1101/2023.03.29.534843

**Authors:** Dafni Skiadopoulou, Jakub Vašíček, Ksenia Kuznetsova, Lukas Käll, Marc Vaudel

## Abstract

Precision medicine focuses on adapting care to the individual profile of patients, e.g. accounting for their unique genetic makeup. Being able to account for the effect of genetic variation on the proteome holds great promises towards this goal. However, identifying the protein products of genetic variation using mass spectrometry has proven very challenging. Here we show that the identification of variant peptides can be improved by the integration of retention time and fragmentation predictors into a unified proteogenomic pipeline. By combining these intrinsic peptide characteristics using the search-engine post-processor Percolator, we demonstrate improved discrimination power between correct and incorrect peptide-spectrum matches. Our results demonstrate that the drop in performance that is induced when expanding a protein sequence database can be compensated, and hence enabling efficient identification of genetic variation products in proteomics data. We anticipate that this enhancement of proteogenomic pipelines can provide a more refined picture of the unique proteome of patients, and thereby contribute to improving patient care.

## Introduction

Genomic variation can affect proteins, their expression, structure [1], degradation rates, or even completely preventing their production [2]. Consequently, cellular functions can be altered, possibly participating in the development of diseases [3]. Therefore, monitoring the proteomic profiles of patients is seen as a promising technique for the development of precision medicine approaches [4]. However, in mass spectrometry-based proteomics, spectra are usually matched to a one-database-fits-all set of protein sequences. Projecting all data onto a database that does not capture the diversity of proteomic samples can yield false positive identifications [5], but more importantly, it creates a bias towards populations of study participants based on their genetic similarity with the reference database.

The personalization of proteomic searches using genomic information is an active field of research in proteogenomics [6]. Typically, proteomic mass spectrometry (MS) data are matched against a database of sequences capturing the products of genomic sequence variation. These databases can be constructed based on genomic or transcriptomic sequencing data, or, when no genomic data are available, using variants from knowledgebases like Ensembl [7]. However, expanding protein sequence databases using sequence variation poses major challenges to the current bioinformatic methods for protein identification: (i) the search space of possible peptides used to match spectra is enlarged, yielding higher processing time and increasing the likelihood of matching a false positive at a given score [8]; and (ii) variant peptides containing the product of an amino acid substitution are highly similar to canonical or modified peptides and thus difficult to confidently identify [9, 10, 11]. These issues, combined with the low sequence coverage of proteomics, make the detection of the products of genetic variation a challenging task, with recent publications showing low identification rates of variant peptides compared to what was expected after analysis at the DNA and RNA level [12, 13]. And when variant peptides are matched to spectra, the evaluation of results remains challenging, often requiring costly experimental validation [12].

The confidence in peptide identification is evaluated by search engines through the matching of the measured spectra with expected fragment ions, and returned as a score. The score is translated as a statistical metric, e.g. a false discovery rate, e-value, or posterior error probability (PEP), after comparison with the estimated null distribution of scores. The reference method for the estimation of the null is the target-decoy strategy, where incorrect sequences, the *decoy* sequences e.g. randomized, shuffled, or reversed, are inserted in the database and compete equally with the sequences of interest, the *target* sequences [14]. However, these scores rely on limited information on the peptides, typically only predicted fragment masses, and usually only consider the most intense peaks in the measured spectra. Bioinformatic approaches were therefore developed that allow re-scoring the matches based on more peptide features, including notably Percolator [15], Scavager [16], and AlphaPept [17]. Notably, the inclusion of predicted retention time [18, 19] and predicted intensities of fragment ions [20, 21, 22, 23] have been demonstrated to increase spectrum identification rates, e.g. with application in immunopeptidomics [24].

In this work we investigate how the inclusion of common germline variation affects the performance of proteomic searches. We demonstrate how variant and canonical peptides distribute in the predicted retention time and fragmentation feature space, and how these can be used to increase the share of confidently identified variant peptides. Together, our results show that with a careful curation of the protein sequence database and using the available tools for post-processing mass spectrometry data, we can gain a better coverage of the variation of the proteome.

## Results

To investigate the influence of including germline variation on the performance of proteomic search engines, we built four protein sequence databases: (i) canonical protein sequences from UniProt [25] as commonly used in proteomic studies, comprising 20,398 distinct protein sequences, (ii) canonical protein sequences plus their isoforms from UniProt, comprising 42,397 distinct protein sequences, (iii) protein sequences from Ensembl [26], featuring all possible isoforms, 92,558 distinct sequences and (iv) protein sequences from Ensembl extended with amino acid substitutions encoded by common genomic variants (minor allele frequency *MAF >* 1%), 155,960 additional distinct sequences, making a grand total of 248,518 distinct sequences. Sequences of common sample contaminants (116 in total) from the cRAP database were also included in all used databases. Then, we searched mass spectrometry data obtained from three samples of healthy tonsil tissue by Wang et al. [12] with a total of 5,085,477 MS2 spectra using X!Tandem [27] against these four databases. The identification results from X!Tandem were then post-processed by Percolator [15] using a set of features proposed in the literature [28]. See methods for details.

### Including germline variation does not impair the identification rate

When germline variation and isoforms are included in the database, there is a substantial increase in the number of sequences that the search engine has to match each spectrum with. In such a case, one would expect a decrease in search performance. However, identification rates at a given false discovery rate (FDR) were nearly identical for the different databases for all three tonsil samples from Wang et al. for both peptide spectrum matches (PSMs) and peptides (Figure 1A and 1B, respectively).

**Figure 1:**
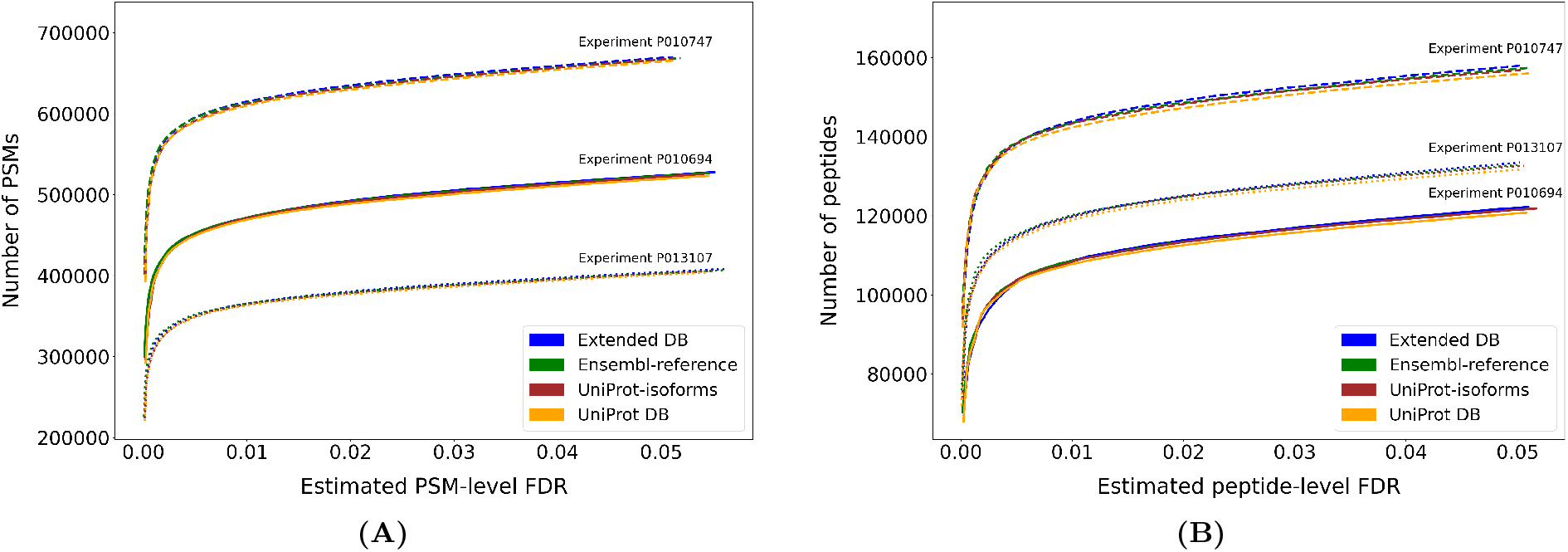
A comparison of the performance when matching spectra to reference databases and extended databases. We searched three different sets of spectra against four different human protein sequence databases. The number of hits obtained at a given FDR threshold is displayed for (A) PSMs and (B) peptide sequences.

The similar or even better performance displayed by the variant-aware Ensembl database indicates that the increase in the number of protein sequences does not create a massive increase in the number of peptides that can match a spectrum. This is also reflected by very similar score distributions for the searches against the canonical UniProt database and the extended Ensembl one (Supplementary Figure S2). The high level of similarity between isoforms and variant proteins might explain that a high number of sequences does not result in a much enlarged search space, in contrast to, e.g., including three-frame translations of untranslated regions (UTRs) or non-coding sections of the genome. Together, these results suggest that extending proteomic sequence databases using common germline variation does not compromise identification rates, while enabling a broader coverage of populations.

### Fragmentation and retention time prediction allows discriminating random matches

Introducing variant sequences however increases the risk of one peptide being difficult to distinguish or even identical to another, possibly from a different protein. The mass difference of an amino acid substitution might even be indistinguishable from a chemical or post-translational modification, yielding equal search scores for the two versions of a peptide that can be encoded by this variant [9]. This further increases the need for a post-search validation of the identifications that can tease apart highly similar peptides. Such tools take advantage of a large number of features to evaluate the quality of PSMs. The features are based on characteristics of the peptide and the spectrum, such as the length of the peptide or the difference between the measured and the theoretic mass over charge ratio. In case two peptidoforms, e.g. a variant and a modified peptide, have the exact same composition, they will be indistinguishable by mass. But these peptidoforms are likely to elute at different retention times, and produce fragments of different intensities. These two characteristics are therefore expected to be a valuable source of information to evaluate the confidence in variant peptide estimations.

For each PSM obtained on the tonsil data by Wang et al. [12], we compared the measured values of retention time and fragmentation with predictions made by DeepLC [19] and MS^2^PIP [21], see methods for details. For peptide retention times, the distances between measured and predicted values were wider for decoys than for target peptides, confirming that excluding PSMs with high deviation in retention time compared to prediction would reduce the prevalence of random matches (Figure 2A, Supplementary Figure S3A). It is important to note that all distributions are centered around zero, which means that there is a substantial number of decoy hits where by chance the retention times expected to be measured for the set of amino acids of these decoy sequences are very close to the measured retention times of the corresponding spectra.

**Figure 2:**
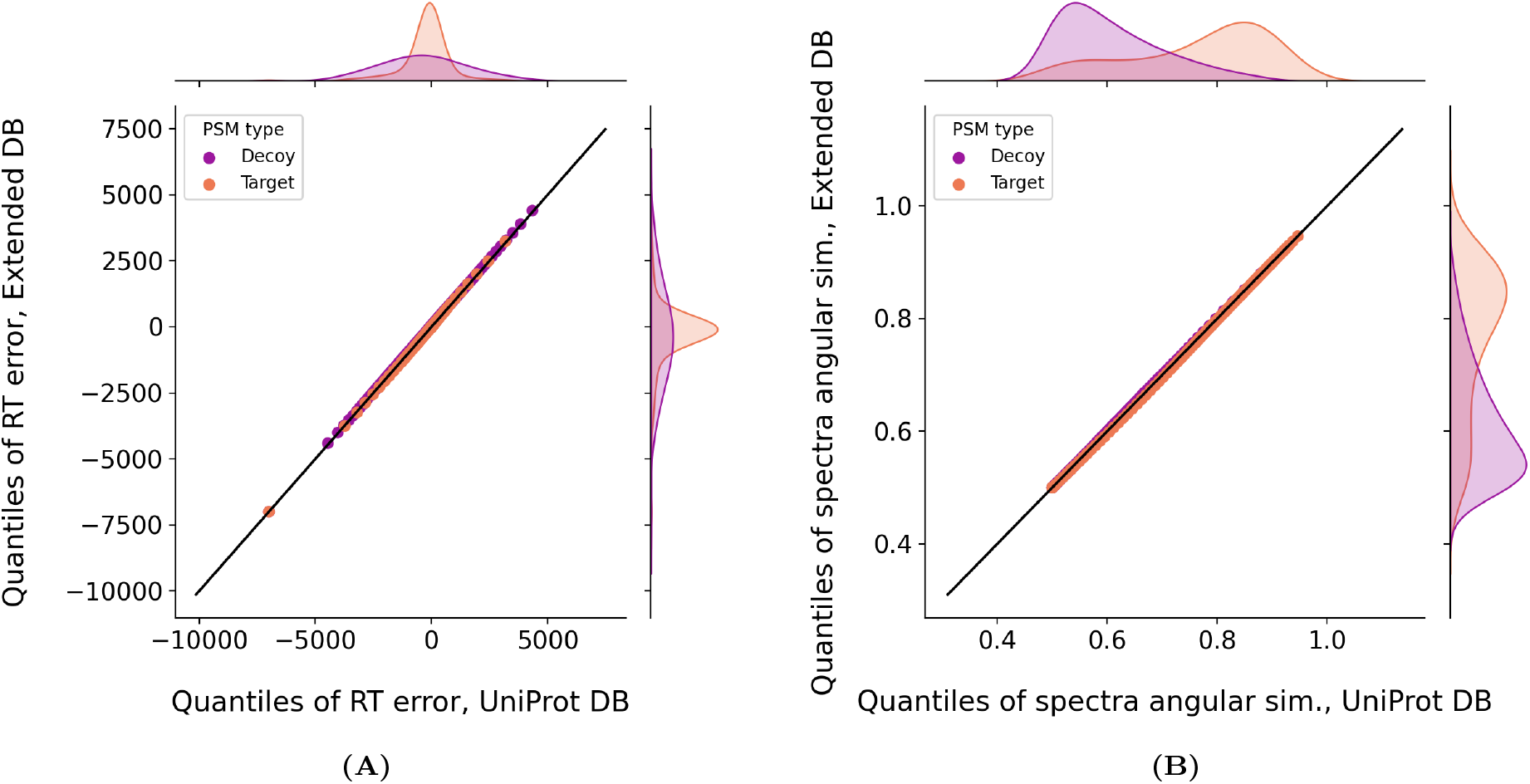
Comparison of PSM features distributions between extended and canonical databases. Q-Q plots that compare the distributions of the retention time error and spectra angular similarity between measured and predicted values of target and decoy hits from the Extended DB and UniProt DB.

For peptide fragment intensities, target hits had a higher share of PSMs with a high similarity between the measured and predicted spectra, and most decoy hits had a very poor agreement between measured and predicted spectra (Figure 2B, Supplementary Figure S3B). This can be explained by the fact that the fragmentation similarity measure compares pairs of matched intensities, in contrast to retention time deviation, which compares two data points. A random match is therefore much less likely to present a good spectrum similarity than a low retention time difference. This indicates that selecting PSMs with high similarity would enrich the data set for high-quality matches. For many PSMs, no measured peak could be matched to predicted peaks, yielding the lowest similarity score (0.5) (81,344 PSMs from the variant-aware Ensembl database: 44,315 targets and 37,029 decoys, 74,582 PSMs from the UniProt database: 41,175 targets and 33,407 decoys). This can be due to a completely wrong match, or to the predictor failing to predict the intensity of some peaks for the given peptide. Given the high prevalence of decoys with very low similarity score, it can be anticipated that most PSMs with low score will be incorrect matches, but one cannot rule out that some good matches will present low similarity scores due to the performance of the fragmentation predictor.

For both retention time and peptide fragmentation, no relevant difference was observed between the different databases (Supplementary Figure S3). The similarity of the distributions of the investigated features for the four sequence databases’ decoy PSMs indicates that there is no obvious bias between the databases. Also, the nearly identical distributions of target PSMs confirms that using the larger databases does not substantially increase the prevalence of hits of lower quality.

When focusing on the results obtained on the variant-aware database, as expected from Figure 2, the decoy hits distribute symmetrically around the zero retention time deviation with most hits at the lowest spectrum similarity, a density of hits decreasing with the similarity, and no relationship is observed between the two features (Figure 3B). On the other hand, the target hits display a similar background of hits supplemented with a dense cloud of PSMs with small retention time deviation and high spectrum similarity, which is likely to contain the best matches. When separating the target PSMs between those mapping to a canonical protein (97.5 %) and those solely mapping to variant peptides (2.5 %), one can see that the variant peptides distribute in this two-dimensional feature space as sampling from the decoy distribution together with a marked cloud of PSMs with low retention time deviation and high spectrum similarity (Figure 3D). This demonstrates that even though the variant peptides present a higher prevalence of PSMs presenting a poor agreement with the predictors than the canonical PSMs, they also have a substantial share of high quality matches. Therefore, the agreement with predictors can help discriminate them from the random matches.

**Figure 3:**
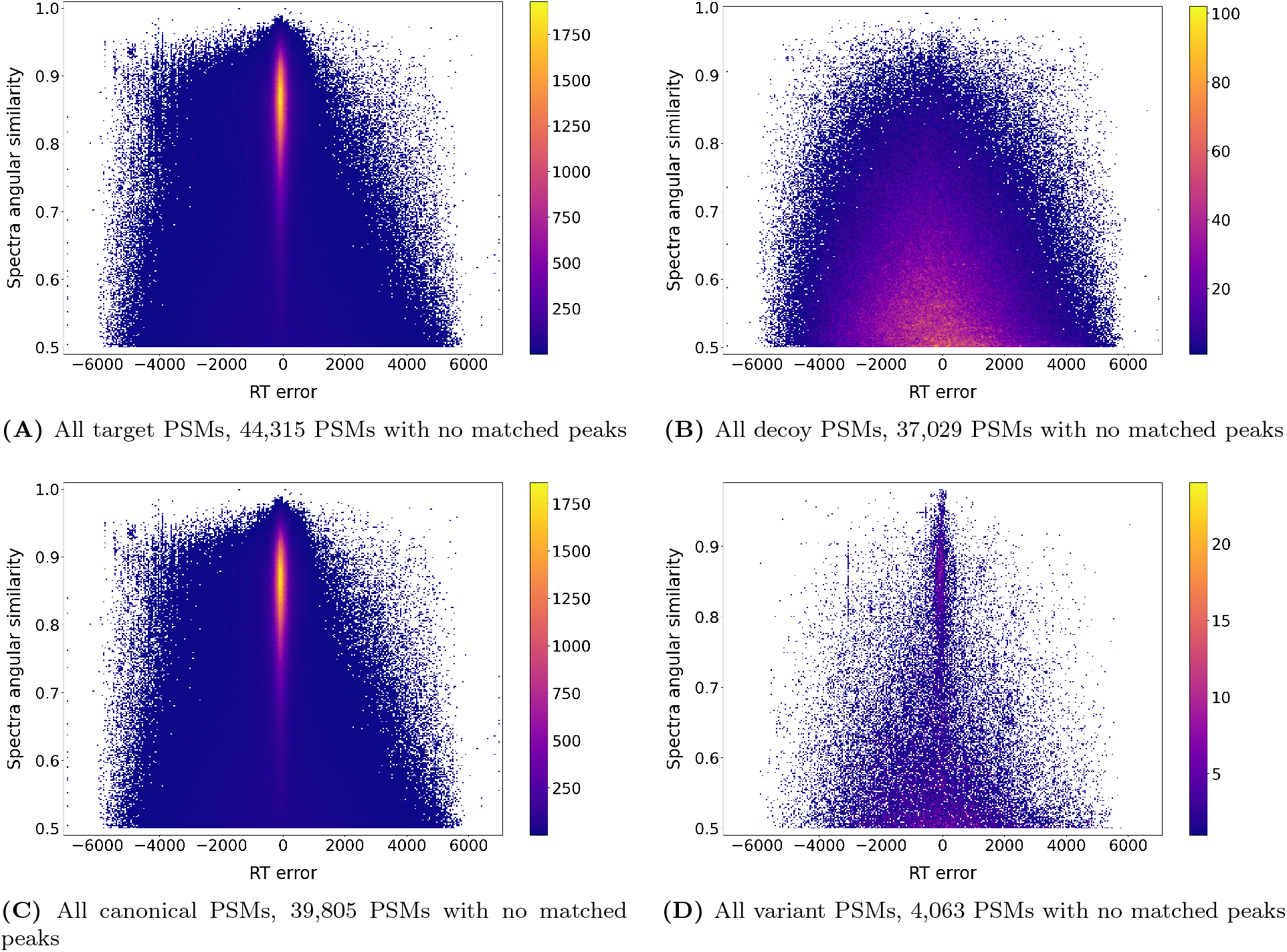
2D-density plot of PSM agreement with retention time and fragmentation predictors. The retention time vs. fragmentation distance to prediction of all (A) target, (B) decoy, (C) canonical, and (D) variant PSMs obtained when searching against the extended protein sequence database. PSMs with no matched peaks are not represented and their prevalence is listed under the plot.

### Percolator combined with predictors increases the identification rate of variant peptide sequences

Since the retention time and fragmentation pattern features capture different aspects of the quality of the match between a spectrum and a theoretical peptide, we hypothesized that their inclusion would enhance the discriminative power of Percolator, and thereby increase the performance when using the database containing the product of germline variation. We therefore extended the set of features given to Percolator to capture the agreement between experimental peptide retention time and fragmentation, and predicted values, making a total of 39 features compared to 18 in the standard set (full list of PSM features available in Materials and Methods). For all three tonsil samples from Wang et al. [12], identification rates were improved when using the extended features (Figure 4). At a global 1 % FDR threshold, Percolator using the new set of features increased the prevalence of PSMs with low retention time and fragmentation deviation from the predicted values, and rejected PSMs with poor retention time or fragmentation pattern matching (Figure 5).

**Figure 4:**
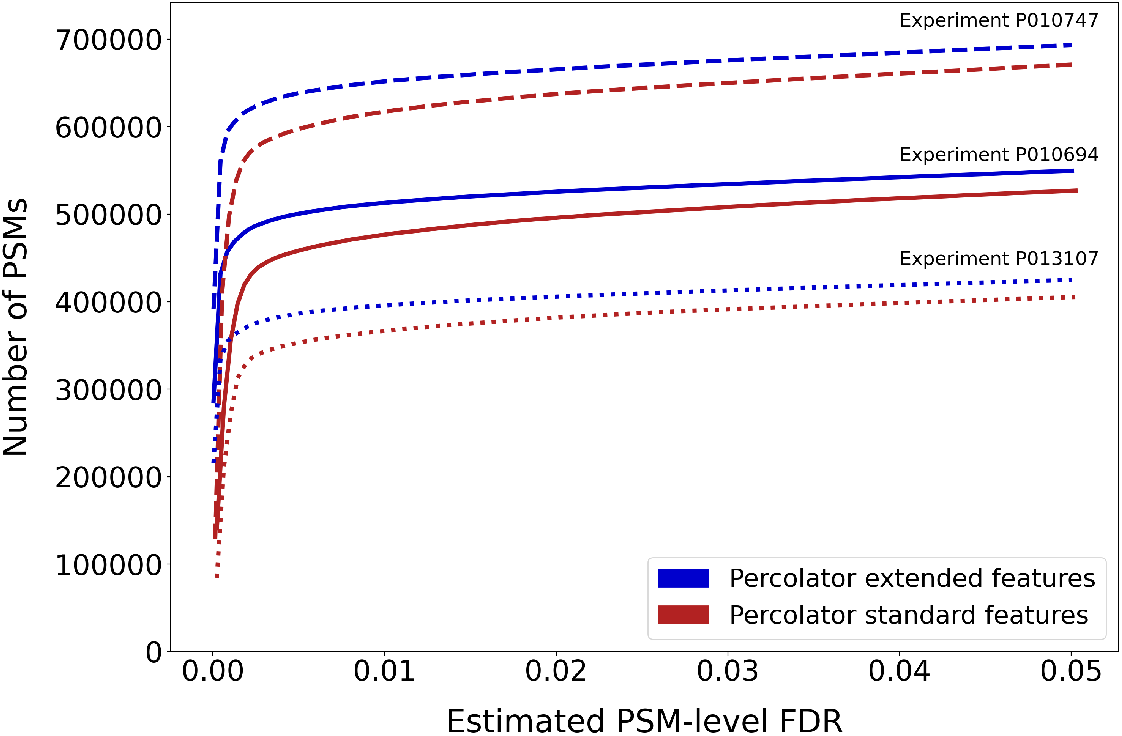
A comparison of the performance of Percolator given the standard and the extended set of features. For all three different sets of spectra that were searched against the extended protein sequences database, the number of PSMs retained at a given FDR threshold is plotted using the standard and extended sets of features for all thresholds up to 5% FDR.

**Figure 5:**
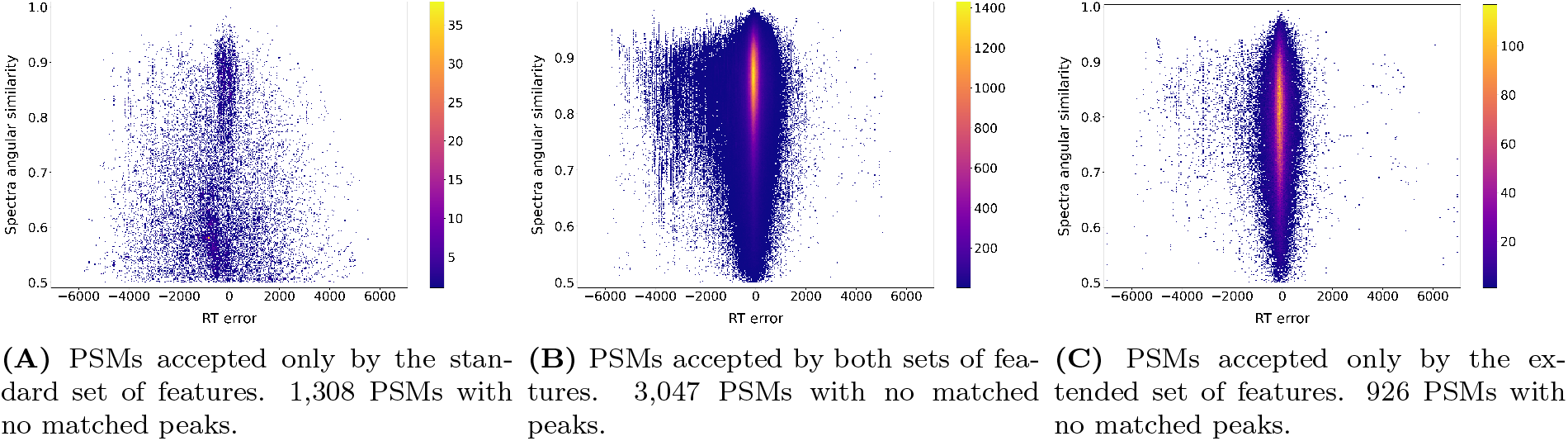
2D-density plot of PSM agreement with retention time and fragmentation predictors for confident PSMs separated based on the set of features supporting their identification. The retention time vs. fragmentation distance to prediction of target peptide-spectrum matches retained at a 1% FDR when Percolator was provided with (A) only the standard set of features, (B) either the standard or extended set of features, and (C) only the extended set of features.

When summarizing the identifications from all three samples, for peptides mapping to a canonical protein sequence, 20,213 PSMs (1.5 %) of the original matches were not retained using the extended features and 115,872 were newly included, representing an increase of 6.9 % (Figure 6, Table 1). When considering distinct peptide sequences, 4,303 sequences (2.2 %) were not retained and 20,759 were newly included, yielding a 8.2 % increase. For variant peptides, 940 PSMs (13 %) were not retained and 1,538 PSMs were newly included, making a 8.3 % increase. When considering distinct peptide sequences, 244 sequences (14.6 %) were not retained and 351 were newly included, making a 6.4 % increase. Thus, using the extended features increased the identification rates for all matches.

**Table 1:**
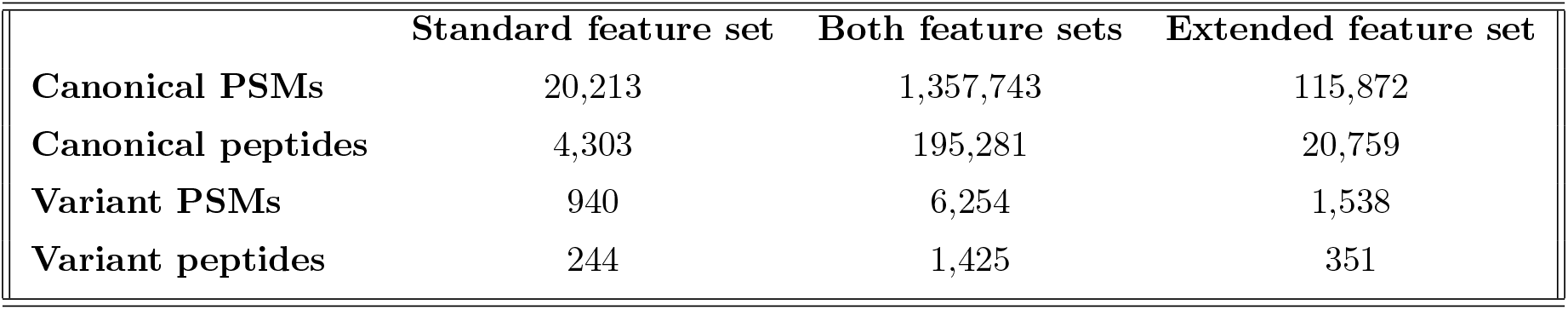
Number of matches retained by Percolator using different sets of features. The number of PSMs and peptide sequences retained using the standard set of features only, either the standard or extended set of features, and the extended set of features only are provided for canonical and variant sequences.

|  | Standard feature set | Both feature sets | Extended feature set |
| --- | --- | --- | --- |
| **Canonical PSMs** | 20,213 | 1,357,743 | 115,872 |
| **Canonical peptides** | 4,303 | 195,281 | 20,759 |
| **Variant PSMs** | 940 | 6,254 | 1,538 |
| **Variant peptides** | 244 | 1,425 | 351 |

**Figure 6:**
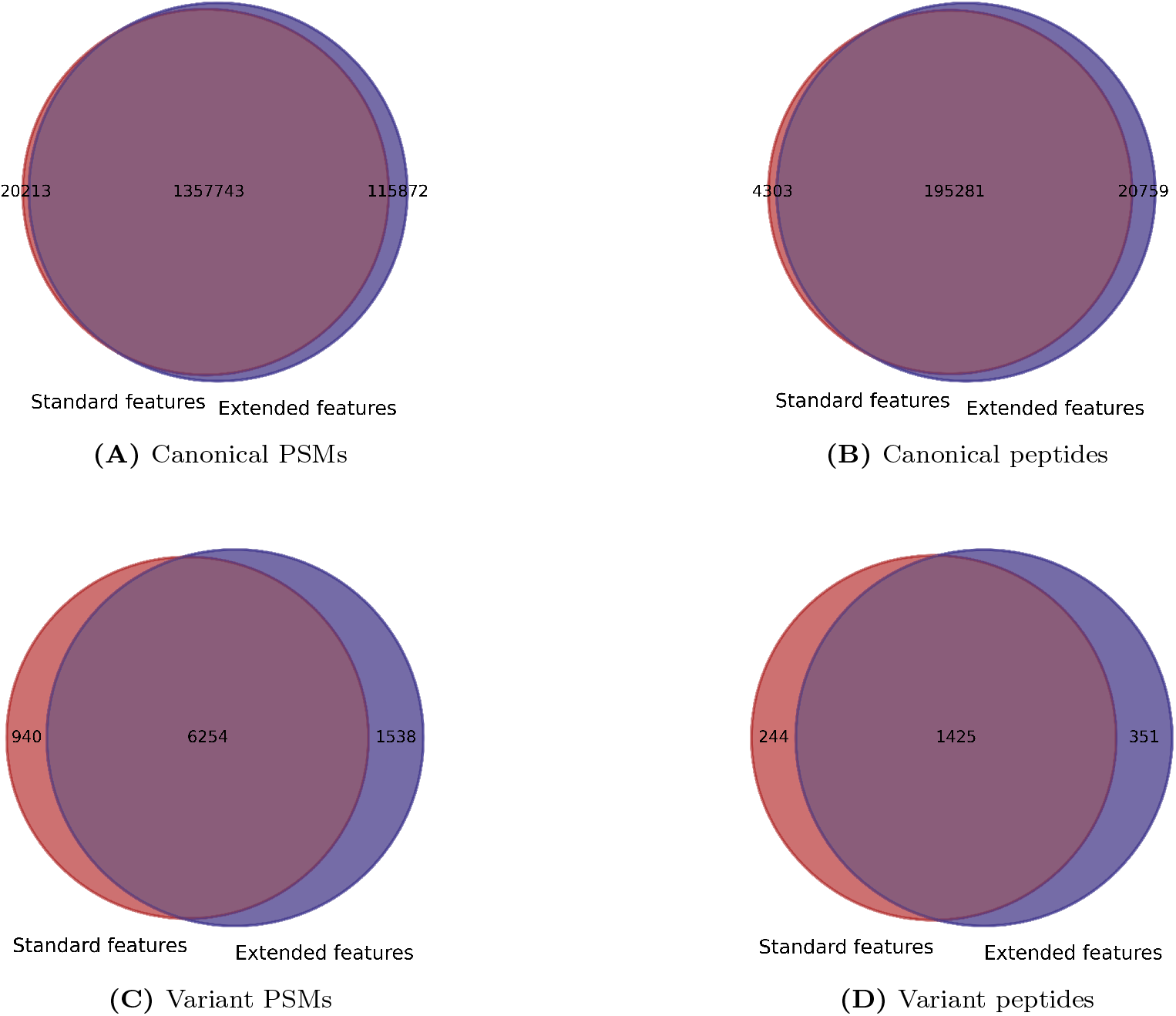
Venn diagrams of the number of PSMs and peptide sequences obtained using different sets of features. The number of PSMs and peptide sequences retained using the standard set of features only, either the standard or extended set of features, and the extended set of features only, are provided for canonical and variant sequences, as listed in Table 1.

Even though the share of variant PSMs and peptide sequences that are gained by the extended set of features is slightly smaller for variant sequences than canonicals, there is a substantially larger percentage of variant PSMs and peptide sequences that are not retained from the standard search. Therefore, the proposed approach manages to eliminate a larger share of random hits mapping to variant sequences from the final confident identifications. Given that variant peptides can be more difficult to distinguish from others, e.g., due to post-translational modification, it is expected that these benefit best from an increased ability to assess the quality of a match. The agreement between PSMs and peptide sequences further indicates that the increase is not only due to the redundant sampling of the same sequence.

The variant PSMs that are not retained using the extended features are mainly the ones that show a large disagreement with the prediction for retention time and/or fragmentation, which makes them less reliable (Figure 7A). While on the other hand, the PSMs that are gained (Figure 7C) together with the ones that are accepted by both sets of features (Figure 7B) display a much better agreement with the predictors: for the majority of them, the retention time of the measured spectrum is very close to the predicted one, and also the intensities of the measured spectrum are highly similar to the predicted fragmentation pattern of the theoretic peptide. Extending the features therefore not only increases the identification rate for variant peptides, it also improves the agreement with predicted retention time and fragmentation.

**Figure 7:**
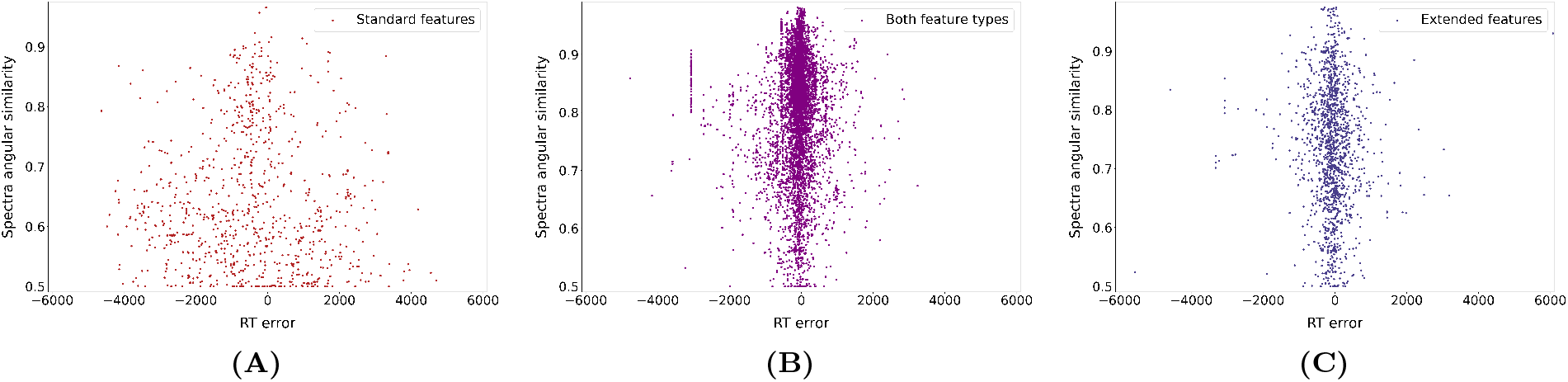
2D-density plot of the variant PSMs accepted using different sets of features. The retention time vs. fragmentation distance to prediction of variant peptide-spectrum matches retained at a 1% FDR when providing Percolator with (A) only the standard set of features, (B) either the standard or extended set of features, and (C) only the extended set of features.

## Conclusion & Discussion

This work focuses on the search for genetic variation products in proteomic data. For this purpose a proteogenomics pipeline was developed and tested on a dataset of healthy tonsil tissue samples available from Wang et al. [12] (Supplementary Figure S1). The search was performed against an Ensembl-based protein database, enriched with the products of common genetic variants and sample contaminants. In order to improve the identification rate for canonical and especially for variant sequences, the retention time and fragmentation pattern of the peptides were used for the computation of additional features for Percolator. The results presented in this paper show that there is indeed a significant influence of these two characteristics in the outcomes of our analysis. By taking them into account, Percolator is able to retrieve a set of accepted PSMs with a greater prevalence of high-quality matches, leading to an increased number of identified peptide sequences.

The performance of peptide identification search engines is strongly affected by the protein sequence database used. In this work the focus was on products of common germline sequence variation which increase the database size but do not yield a search space explosion compared to rare or somatic mutations. If rare (*MAF* ≤ 1%) or somatic variants were also included, then for most of the proteins there would be orders of magnitude more unique sequences that would need to be included in the extended database. Similarly, if the search were also aiming at the products of untranslated regions or non-coding variants, then the massive size of the resulting database would pose several challenges and the prevalence of false positive hits would increase significantly. In these cases, the improvement of Percolator’s evaluation with the additional features of retention time and fragmentation pattern are likely to also have a positive influence on the performance of a proteogenomics pipeline.

Our study focused solely on the identification of PSMs and peptide sequences. Accounting for common germline variation remains to be integrated with PTM detection and localization methods to enable the identification of peptides. Similarly, new methods and tools need to be developed to consolidate the variant-aware peptide information at the gene or protein level. But overall our results support that current proteomic pipelines have the potential to account for products of germline genetic variants. Routinely including genetic variation in proteomic analyses holds the promise to increase their value in medical and population studies, and especially in precision medicine approaches. It also provides a simple alternative to projecting all data onto an arbitrary reference genome, hence enabling a better and fairer coverage of populations.

## Materials and Methods

### Data samples

The processed samples were published by Wang et al. [12] and downloaded from the PRIDE repository with identifier PXD010154. From this dataset the chosen subset of samples consists of 106 mass spectrometry raw files of healthy tonsil tissues acquired from 3 different experiments with identifiers P010747, P010694, and P013107. Briefly, the proteins were digested with trypsin and analyzed by tandem mass spectrometry coupled with liquid chromatography (LC-MS/MS) using a Q Exactive Plus mass spectrometer, yielding 5,085,477 MS/MS spectra (Exp. P010747: 1,834,613 MS/MS spectra, Exp. P010694: 1,695,460 MS/MS spectra, Exp. P013107: 1,555,404 MS/MS spectra). For more details on the data generation please refer to the original publication by Wang et al. [12].

### Protein databases

The search was done against four different protein databases, three that included the canonical human proteome and/or protein isoforms and one that also included genetic variation products. The three canonical databases were: (i) the homo sapiens complement of the UniProtKB database downloaded on September 20th 2022 (20,398 distinct protein sequences), (ii) the homo sapiens complement of the UniProtKB database including protein isoforms downloaded on January 9th 2023 (42,397 distinct protein sequences), and (iii) the canonical database of protein isoforms of homo sapiens taken from Ensembl v.104 (92,558 distinct sequences).

The extended database included the protein products of genetic variants, appended with the canonical database of protein isoforms taken from Ensembl v.104 (248,518 distinct sequences). We included variants with minor allele frequency *>* 1%, taken from Ensembl v.104, and six-frame translations of variant cDNA were obtained using the Python tool py-pgatk [7]. We then included only translations of the main open reading frame (mORF) in each transcript, as annotated per Ensembl v.104. Translations of cDNA without an annotated mORF were not included in the database. Decoy sequences were generated using the algorithm DecoyPYrat [31], implemented by py-pgatk.

All databases were supplemented with sample contaminants from the common Repository of Adventitious Proteins (cRAP, thegpm.org/crap).

### Proteomic search

The RAW files were converted to mzML files using ThermoRawFileParser version 1.3.4 [32]. The mzML files were searched using the X!Tandem search engine [27] operated through the SearchGUI interface version 4.0.41 [33]. Search settings were: (1) specific cleavage by trypsin with a maximum number of 2 missed cleavages; (2) Carbamidomethylation of C as fixed and Oxidation of M, Deamidation of N and Q, and Acetylation of protein N-terminus as variable modifications; (3) peptide maximum length of 40 amino acids; (4) precursor and fragment ion tolerance of 10 ppm. The refinement step of X!Tandem was disabled. PeptideShaker version 2.2.9 [34] was used to process the output of X!Tandem and generate standardized PSM exports.

### Peptide feature predictors and confidence scoring

Percolator version 3.5 [15] was used for the statistical evaluation of the resulting peptide-to-spectrum matches (PSMs). For each PSM, a set of features commonly used for Percolator [28] was generated using PeptideShaker, referred to as the *standard* set of features, and described in Table 2. This standard set of features was extended with novel features capturing the agreement between PSMs and predicted retention times and fragmentation, resulting in a new set of features referred to as the *extended* set of features, and described in Table 3.

**Table 2:** Standard set of features representing PSMs.

|  |  |  |
| --- | --- | --- |
| 1 | measured_mz | The measured mass over charge ratio |
| 2 | mz_error | The difference between theoretic and measured mass over charge ratio |
| 3 | pep | Posterior error probability estimate based on the search engine score provided by PeptideShaker |
| 4 | delta_pep | Difference in posterior error probability estimate between the peptide candidate and the next best scoring peptide candidate for this spectrum |
| 5 | ion_fraction | The fraction of matched b and y ions |
| 6 | peptide_length | The length of the matched peptide, in number of amino acids |
| 7-9 | charge_{2-4} | One-hot encoding of the charge state |
| 10-14 | isotope_{0-4} | One-hot encoding of the isotopic difference of the measured mass over charge ratio |
| 15-18 | {unspecific/enzymatic_[N,C]/enzymatic} | One-hot encoding of the peptide enzymaticity |

**Table 3:** Extended set of features representing PSMs.

|  |  |  |
| --- | --- | --- |
| 1-18 | "Standard features" | PSM features of the standard set presented at Table 1 |
| 19 | measured_rt | The measured retention time |
| 20 | rt_Abs_error | The absolute distance of measured and predicted retention time |
| 21 | rt_Square_error | The squared distance of measured and predicted retention time |
| 22 | rt_Log_error | The logarithmic distance of measured and predicted retention time |
| 23 | matched_peaks | The percentage of predicted peaks matched with an observed peak |
| 24 | spectra_log | The logarithmic distance of the observed and predicted spectrum |
| 25 | spectra_cos_similarity | The cosine similarity of the observed and predicted spectrum |
| 26 | spectra_angular_similarity | The angular similarity of the observed and predicted spectrum |
| 27 | spectra_cross_entropy | The cross entropy of the observed and predicted spectrum |
| 28 | b_ion_coverage | Number of consecutive amino acids covered by b ions starting from the N-terminus |
| 29-33 | "Fragmentation features for b ions" | Features 23-27 calculated on the predicted b ions only. |
| 34 | y_ion_coverage | Number of consecutive amino acids covered by y ions starting from the C-terminus |
| 35-39 | "Fragmentation features for y ions" | Features 23-27 calculated on the predicted y ions only. |

DeepLC version 1.0.0 [19] was used to compute predictions of the retention time of each theoretic peptide of each PSM. The retention times of the confident (q-value ≤ 0.01) PSMs according to Percolator using the standard features were used to calibrate the predictions of DeepLC. This was then compared with the retention time reported for the matched spectrum and three different metrics (absolute distance, square distance, and logarithmic distance) were calculated and used as PSM features. The retention time error used in the figures of the Results section correspond to the residuals of a linear regression model computed on the measured and predicted retention times of the confident target hits from Percolator run using the standard set of features.

The peptide fragmentation predictions were obtained from MS^2^PIP version 3.6.3 [21] and were compared against the peaks of the experimental spectrum after matching of the predicted and observed fragment peaks using a 10 ppm threshold and the normalization of the intensities of the peaks. The features used to evaluate the concordance between experimental and predicted spectra were: (1) the percentage of predicted peaks matched with an observed one, (2) the logarithmic distance, (3) the cosine and angular similarity and (4) the cross entropy between the spectra. These features were calculated when taking into account the predicted b and y ions separately and also with all the ions combined. In addition, the number of consecutive amino acids matched from the N and C termini were computed for the b and y ions, respectively.

## Code availability

All the steps of the proteogenomic pipeline described above are implemented in a Snakemake [35] workflow (version 6.8.0). The post-processing of the results of Percolator and the creation of the figures were conducted using custom scripts available at the GitHub repository (https://github.com/ProGenNo/VariantPeptideIdentification). A list with the required software packages together with further documentation and links to supplementary data are also included in that GitHub repository.

## Acknowledgements

This work was supported by the Research Council of Norway (project #301178 to MV) and the University of Bergen, and by the Swedish Research Council (grant 2017-04030 to LK).

This research was funded, in whole or in part, by the Research Council of Norway 301178. A CC BY or equivalent license is applied to any Author Accepted Manuscript (AAM) version arising from this submission, in accordance with the grant’s open access conditions.

