## Supplementary material, additional figures. for "Retention time and fragmentation predictors increase confidence in variant peptide identification"

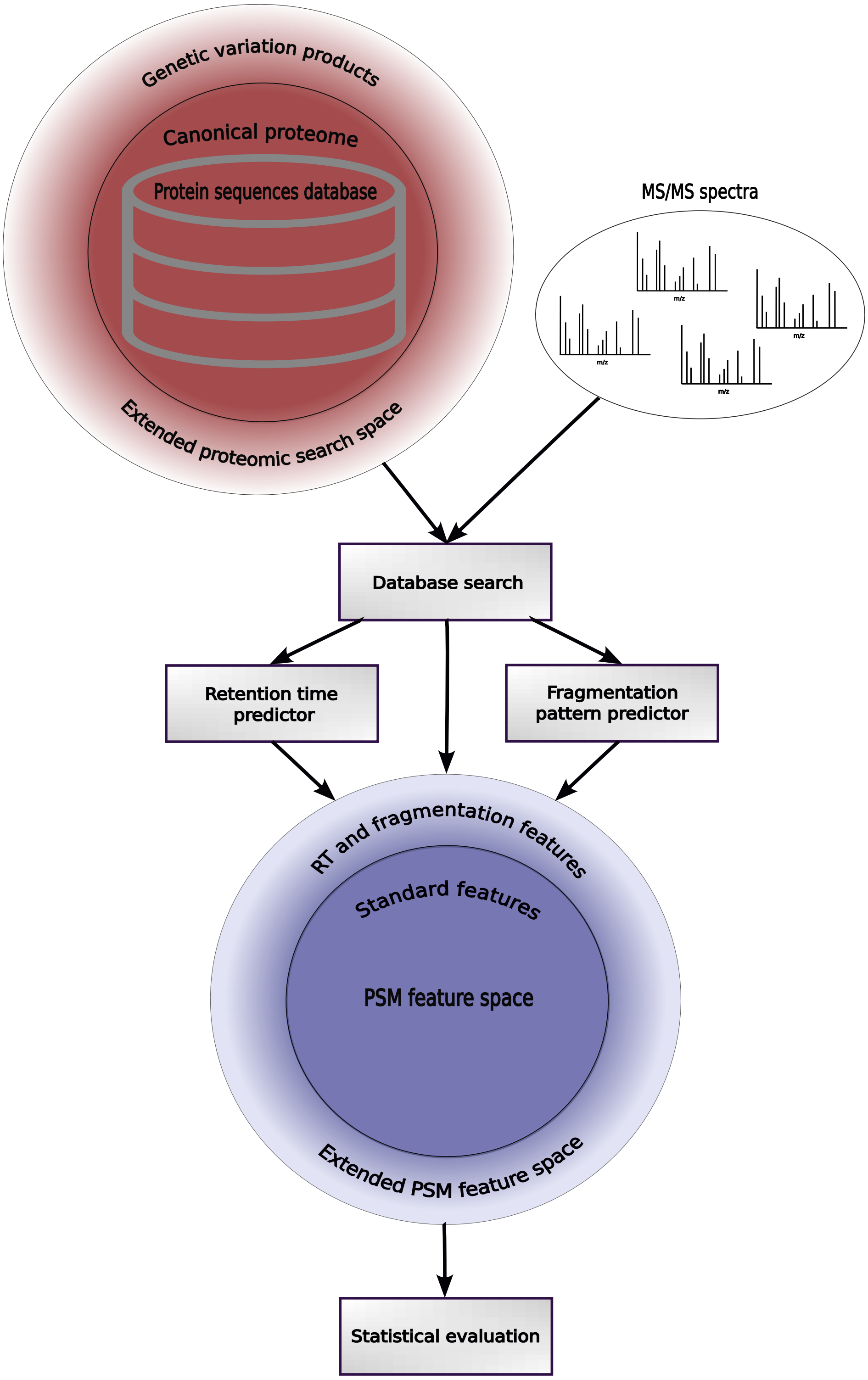


**Figure S1:** Proposed proteogenomics pipeline.

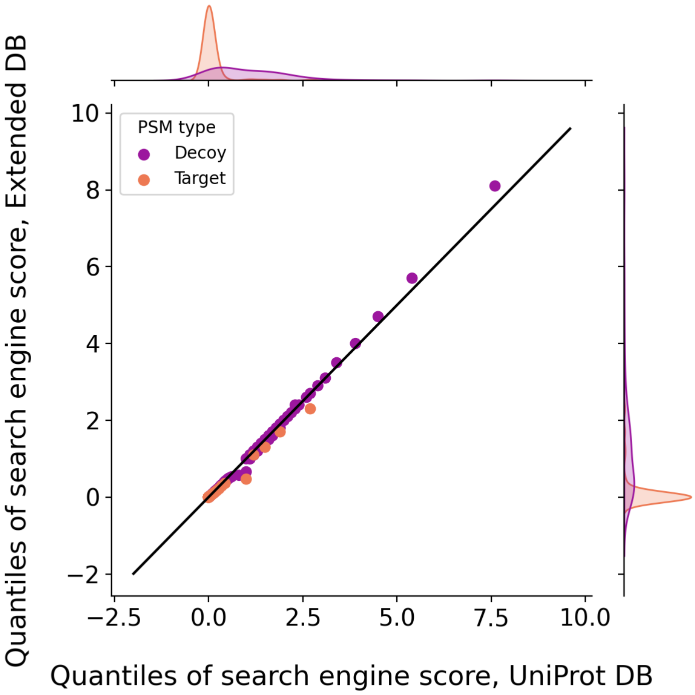


**Figure S2: Comparison of search engine score distributions between extended and canonical databases.** Q-Q plots that compare the distributions of the search engine score of target and decoy PSMs at a 5% FDR from the variant-aware Ensembl database and UniProt DB.

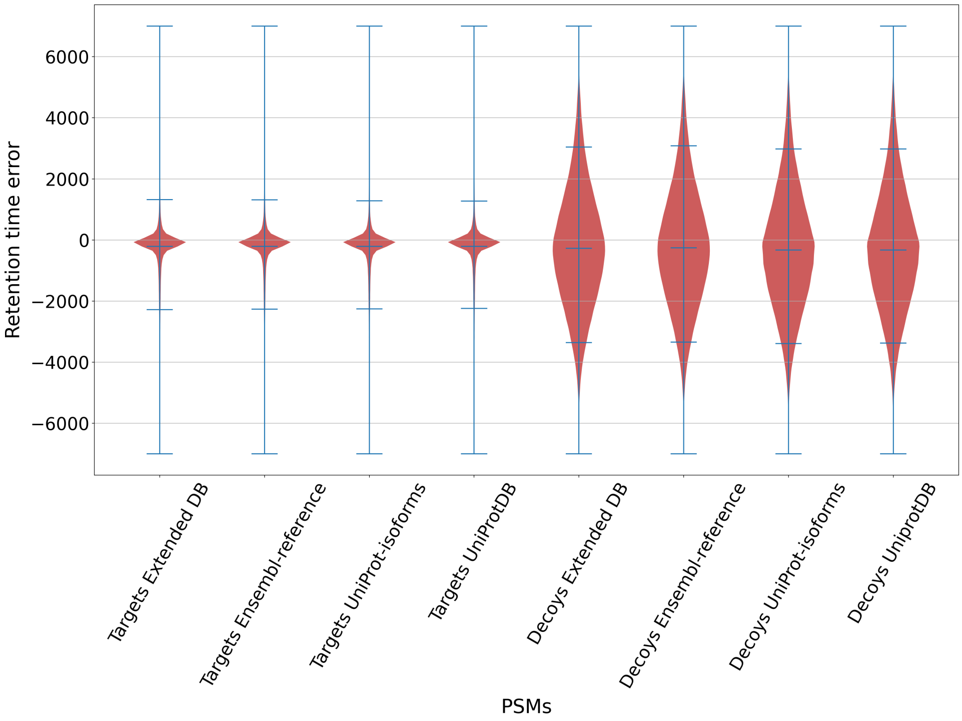


**(A)**

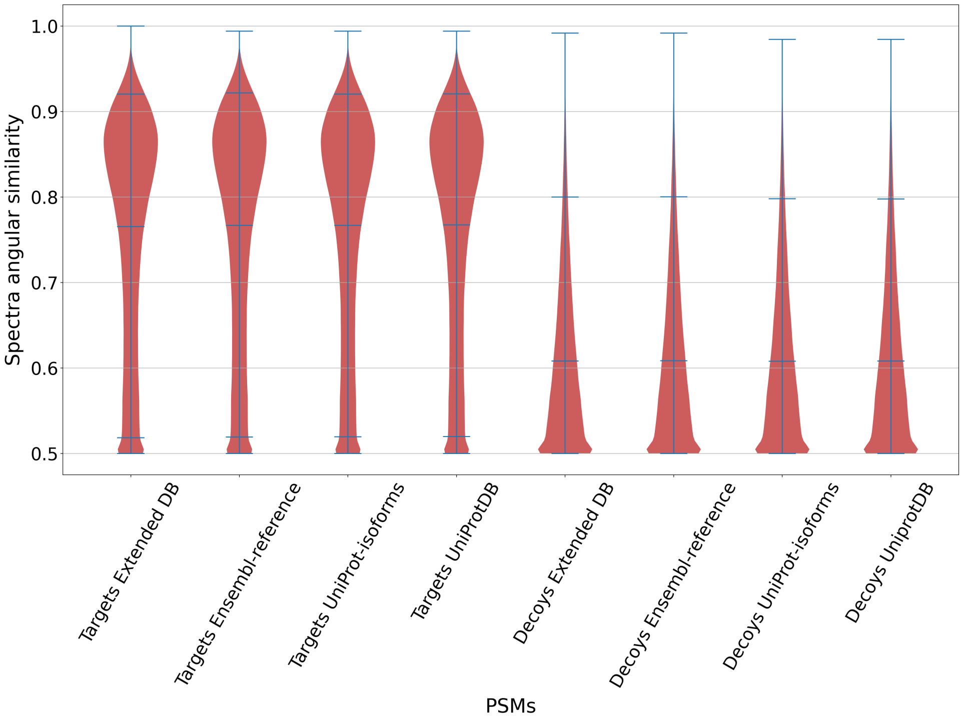


**(B)**

**Figure S3: Violin plots of PSM features obtained from the search against the four protein databases for target and decoy hits.** The two features presented in these figures represent the agreement of the matches with peptide (A) retention time and (B) fragmentation obtained using DeepLC and $MS^2$PIP, respectively. See methods for details on how these features are computed.
